# Encoding of 3D head direction information in the human brain

**DOI:** 10.1101/335976

**Authors:** Misun Kim, Eleanor A. Maguire

**Author notes:** Grant sponsors: Wellcome (101759/Z/13/Z awarded to E.A.M.; 203147/Z/16/Z awarded to the Centre; 102263/Z/13/Z awarded to M.K.); Samsung Scholarship (awarded to M.K.). Corresponding author: Eleanor Maguire Wellcome, Centre for Human Neuroimaging, Institute of Neurology, University College, London, 12 Queen Square, London WC1N 3AR, UK; E.

## Abstract

Head direction cells are critical for navigation because they convey information about which direction an animal is facing within an environment. To date, most studies on head direction encoding have been conducted on a horizontal two-dimensional (2D) plane, and little is known about how three-dimensional (3D) direction information is encoded in the brain despite humans and other animals living in a 3D world. Here, we investigated head direction encoding in the human brain while participants moved within a virtual 3D “spaceship” environment. Movement was not constrained to planes and instead participants could move along all three axes in volumetric space as if in zero gravity. Using functional MRI multivoxel pattern similarity analysis, we found evidence that the thalamus, particularly the anterior portion, and the subiculum encoded the horizontal component of 3D head direction (azimuth). In contrast, the retrosplenial cortex was significantly more sensitive to the vertical direction (pitch) than to the azimuth. Our results also indicated that vertical direction information in the retrosplenial cortex was significantly correlated with behavioural performance during a direction judgment task. Our findings represent the first evidence showing that the ‘classic’ head direction system that has been identified on a horizontal 2D plane also seems to encode vertical and horizontal heading in 3D space in the human brain.

## 1. INTRODUCTION

Knowing one’s orientation within an environment is critical for navigation. Head direction (HD) cells in a network of brain structures including anterior thalamus, presubiculum, retrosplenial cortex (RSC) and entorhinal cortex (EC) are typically regarded as comprising the ‘neural compass’ because they fire when an animal is facing in a particular direction in space (for a recent review, see Cullen & Taube, 2017). HD cells have been mostly observed in rodents (Taube, 2007), but also in primates (Robertson et al., 1999), while human neuroimaging studies have detected HD information in relevant brain structures (Baumann & Mattingley, 2010; Marchette et al., 2014; Chadwick et al., 2015; Shine et al., 2016). HD cells integrate multisensory information (vestibular, visual, proprioceptive) to update an animal’s heading, and this direction information is critical for maintenance and updating of the spatial map of an environment that is encoded by place cells and grid cells (Calton et al., 2003; Burak & Fiete, 2009). Thus, cells encoding 3D direction information would be crucial for navigation in the three-dimensional (3D) world in which we live. However, most studies of HD encoding have been conducted on a horizontal two-dimensional (2D) plane and there is a dearth of knowledge about how 3D direction information is encoded in the brain.

An early study observed a small number of vertical pitch-sensitive cells in the lateral mammillary nuclei of rats that could potentially be involved in 3D direction encoding (Stackman & Taube, 1998). However, these cells responded only when a rat was looking up almost 90°. The absence of cells tuned to an intermediate angle, and limitations in the apparatus which could not unambiguously detect pitch angles smaller than 40°, made it difficult to provide clear evidence of vertical direction encoding. In several other studies, HD cells were recorded when rats were climbing a vertical plane or were on a ceiling (Taube et al., 2004; Calton & Taube, 2005; Taube et al., 2013). The results indicated that HD cells responded to an animal’s direction relative to the local plane of locomotion, as if the new vertical plane was an extension of the horizontal floor. More recently, Page et al. (2018) proposed a dual-axis rotation rule for updating HD cells based on the HD cells’ responses when a rat moves between multiple planes. These studies have significantly extended our understanding of HD cells by incorporating multiple interconnected planes within a 3D world. However, movements are not always restricted on planes. Primates, who are evolutionally closer to humans than rodents, explore volumetric spaces like arboretums. Human astronauts, pilots and divers also have complete degrees of freedom in 3D space. Although flying and underwater movement are less common forms of behaviour in humans, they nevertheless occur. Therefore, the question of how this is accomplished, and whether humans possess mental representations of volumetric 3D space and can process 3D HD signals, is important to understand.

A recent breakthrough in the study of 3D HD arose from bats (Finkelstein et al., 2015). HD cells were recorded in the bat presubiculum in multiple environments ‐ a horizontal 2D plane, a vertical ring platform and a 3D arena. A large portion of cells were sensitive to azimuth (horizontal direction) only, but a significant number of cells were tuned to various vertical pitches (unlike the rat lateral mammillary cells, which only responded to extreme tilt in Stackman & Taube, 1997) or to 3D direction (“pitch x azimuth conjunctive cells”). An interesting anatomical gradient was also observed in that pure azimuth cells were more abundant in the anterolateral part of presubiculum, whereas pure pitch and conjunctive cells were more numerous in the posteromedial part of presubiculum. These findings provide strong evidence that 3D direction information is present in the bat presubiculum which could be used to generate a mental map of 3D space. In humans, a few functional MRI (fMRI) studies have investigated the neural correlates of vertical heading (Indovina et al., 2013; Indovina et al., 2016; Kim et al., 2017), but in these studies, participants were constrained to a rollercoaster-like track and so the neural basis of complete 3D directional encoding remains unknown.

In the present study, we used an fMRI multivoxel pattern similarity analysis to investigate how 3D direction information was encoded in the human brain when participants explored a volumetric 3D virtual environment, where their movements were not restricted to tracks or planes, as if they were flying in zero gravity. We believe that this unconstrained 3D movement was the most appropriate setup for testing 3D HD encoding, even though such flying is a less common behaviour for most humans. One could also study 3D head tilt by using reaching or grasping behaviour. However, this egocentric representation of 3D space is not of primary interest here, rather we were concerned with understanding allocentric representations in 3D. Our main goal was to test whether vertical and horizontal direction information was encoded using the well-established system known to be involved in supporting HD encoding in 2D navigation, namely the thalamus, subiculum, RSC and EC.

## 2. MATERIALS AND METHODS

Aspects of the methods have been reported previously in our study which investigated grid cells in 3D space using the same fMRI dataset (https://www.biorxiv.org/content/early/2018/03/14/282327), and are reprised here for the reader’s convenience. Of note, the analyses of vertical and horizontal direction encoding in 3D space reported here are completely original and have not been published elsewhere.

### 2.1 Participants

Thirty healthy adults took part in the experiment (16 females; mean age = 25.9 ± 4.8 years; range 19-36 years; all right-handed). All had normal or corrected-to-normal vision and gave informed written consent to participation in accordance with the local research ethics committee.

### 2.2 The virtual environment

The virtual environment was composed of two distinctive rectangular compartments, called here room A and room B for convenience, which were linked by a corridor (Figure 1a). Participants were instructed that they were inside a virtual zero gravity “spaceship” where they could move up, down, forwards and backwards freely. The walls, floors and ceilings had different textures which provided orientation cues. Snapshots of the virtual environment as seen from a participant’s perspective during scanning are shown in Figure 1b-e. The virtual environment was implemented using Unity 5.4 (Unity Technologies, CA, United States) with textures and sci-fi objects downloaded from the Unity Asset Store. The virtual environment can be viewed at: www.fil.ion.ucl.ac.uk/Maguire/spaceship3D.

**Figure 1.**
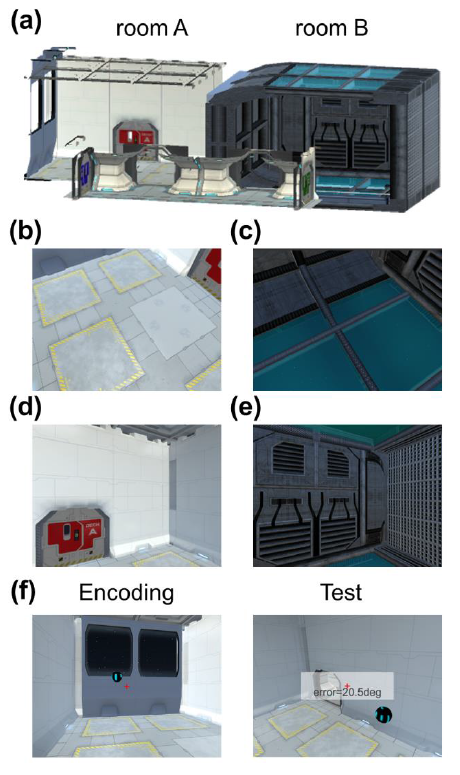
The virtual environment and the pre-scan task. (a) An overview of the virtual spaceship composed of two rooms linked by a corridor. Some walls are shown as transparent here for display purposes. (b-e) Example views from a participant’s perspective during scanning. (b) and (c) are views when a participant is facing down in room A and room B, respectively. (d) and (e) are views when a participant is facing straight ahead in room A and room B, respectively. (f) In a pre-scan task, participants pointed towards the remembered locations of balls while positioned at random locations and then they received a feedback on their decision in terms of angular deviation. Of note, participants performed this task while wearing a VR head-mounted display, which has a wider field-of-view and stereoscopic vision, therefore the example pictures shown here are approximate to the actual views experienced by participants.

The virtual spaceship was rendered on two different mediums for pre-scanning tasks and scanning tasks respectively: a head-mounted virtual reality (VR) display (Samsung Gear VR, model: SM-R322 with a Samsung Galaxy S6 phone) and a standard computer screen (Dell Optiplex 980 with an integrated graphic chipset).

The head-mounted display provided participants with a fully immersive sensation of 3D space via its head motion tracking system, stereoscopic vision and wide field-of-view (96°). A rotation movement in the VR display was made by a participant’s physical head rotation and a forward/backward translational movement was made by a button press on the Bluetooth controller (SteelSeries Stratus XL, Denmark). For example, a participant could move up to the ceiling in the virtual spaceship by physically looking up and pressing the forward button on the controller. To rotate to the right, they physically rotated their head to the right or rotated their whole body when the required rotation was beyond the range of neck rotation. Participants could only move in parallel to their facing direction, and not straight up or down, or side to side. This allowed us to avoid a discrepancy between the HD and movement direction, because this can confound responses in HD cells (Raudies et al., 2015). For ease of rotation, participants were seated on a swivel chair throughout. The VR display was used to provide multisensory (visual, vestibular and proprioceptive) inputs to the HD system. A previous study (Shine et al., 2016) suggested that exposure to both visual and vestibular stimuli during the pre-scan period with a VR head-mounted display might lead to a recapitulation of body-based information during later fMRI scanning, where only visual input is available due to head immobilisation. This pre-exposure to vestibular cues could be particularly important for detecting heading signals in thalamus (Shine et al., 2016). Of note, in our study and that of Shine et al. (2016), head rotation stimulated the semicircular canals in vestibular system, however, linear acceleration signals, which stimulate the otoliths, were absent because participants made virtual translation movements using a controller.

During fMRI scanning, participants watched a video that was rendered on a standard computer screen (aspect ratio = 4:3). The video was a first-person perspective that gave the participants the feeling of moving in the virtual spaceship (details of the tasks are provided in the next section). The stimuli were projected on the screen using a projector at the back of the MRI scanner bore (Epson EH-TW5900 projector), and participants saw the screen through a mirror attached to the head coil. The screen covered a field of view of ~19° horizontally and ~14° vertically.

### 2.3 Tasks and procedure

#### 2.3.1 Pre-scan: familiarisation

Participants first familiarised themselves with the VR head-mounted display and the controller during a simple “ball collection” task (duration = 5 min). Multiple balls were scattered in the spaceship and participants moved to the balls one by one. When they arrived at a ball, they received auditory feedback (a ‘ping’ sound). The primary purpose of this task was to familiarise participants with controlling their movements in the virtual environment via head/body rotations and button presses on the controller. In addition, participants were asked to pay attention to the overall layout of environment for later tasks. This ball collection task also ensured that the participants visited every part of the virtual environment.

#### 2.3.2 Pre-scan: pointing task

After the initial familiarisation period, participants performed a spatial memory task which required a good sense of direction in the virtual 3D spaceship (duration = 15 ± 2 min, Figure 1f). While wearing the head-mounted display, at the beginning of each trial, participants were placed in one of the two rooms in the spaceship. There was one floating ball in the room and participants had to memorise the location of the ball. During this encoding phase (duration = 18 s), participants could move freely and they were instructed to look at the ball from various directions and distances in order to learn the precise location of the ball. The ball then became invisible and a participant was transported to a random location. Participants were then required to look towards the remembered location of the ball and press a button when they had made their decision, after which feedback was provided in the form of the absolute 3D angular deviation from the true direction (Figure 1f). Throughout the task (encoding and testing) a small red crosshair was shown to aid orientation (Figure 1f).

In the majority of trials (“within-room”, n = 16), testing took place in the same room where the ball was located during encoding. There were six additional trials where testing occurred in the other room; for example, participants encoded the ball’s location in room A but they were placed in room B during the test phase, requiring them to point to the ball behind the wall. These “across-room” trials were included in order to encourage participants to build an integrated map of the whole spaceship that was not limited to a local room. An integrated mental representation was important for the later fMRI analyses because we searched for direction information that was generalised across the two rooms.

#### 2.3.3 Scanning: direction judgment task

During scanning, participants watched a video rendered on a standard display and performed a direction judgment task. The video provided participants with the feeling that they were flying in a controlled 3D trajectory within the spaceship (Figure 2a; see also Supporting Figure S1). Similar to the pre-scan task, participants were moved in parallel to their heading direction (e.g. they were tilted up when they moved upwards). The pre-programmed video allowed tight control of location, direction and timing for all participants. The trajectory consisted of multiple short linear movements (each of 3 s, and this was the period included in the fMRI analysis, see Section. 2.6.2) followed by rotation (2/2.6 s). Ideally, we would have sampled all possible directions in 3D space (from ‐180° to 180° horizontally and from ‐90° to 90° vertically), but we restricted the range of linear movement directions in order to acquire reliable measurements of the neural responses to each direction within a reasonable scanning time.

**Figure 2.**
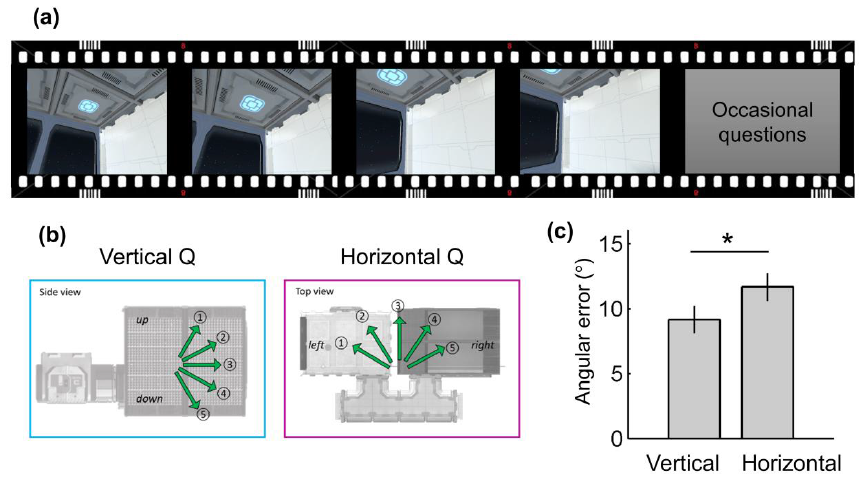
The direction judgment task during scanning. (a) Participants watched a video that provided the sensation that they were moving inside a virtual spaceship. (b) Occasionally, participants were asked to indicate either the vertical or horizontal direction of their last movement. (c) They were more accurate at answering vertical than horizontal questions. Error bars are SEM adjusted for a within-subjects design (Morey, 2008). *p = 0.02

We sampled five levels of horizontal azimuth and five levels of vertical pitch from ‐60° to 60° with 30° steps, resulting in 25 unique 3D directions (Figure 2b). A smooth trajectory was used without abrupt rotations (e.g. if a participant’s previous direction was 0°, the participant would be facing 0 ± 30° after a turn). A constant linear and angular velocity was applied in order to control the velocity, which can modulate the firing rate of HD cells (Stackman & Taube, 1998). If a participant reached the boundary of the spaceship, a blank screen appeared for two seconds and then the next trajectory started from the other end of the spaceship. Twenty five percent of the time, a question screen appeared immediately after a linear movement, and participants indicated the direction of their last movement by pressing an MR-compatible button pad (a 5-alternative forced choice question with a time limit of 5 s, mean response time = 1.7 ± 0.4 s, Figure 2b). This direction judgment task ensured participants kept track of their movements during scanning. They received feedback after each response; the correct direction was shown on the screen if they chose the wrong direction. Since vertical or horizontal direction questions were randomly presented, participants were required to know their 3D direction throughout. Note, this occasional direction judgement was not included in the time period used to estimate neural responses to 3D directions in the main fMRI analysis. The two compartments of the spaceship were visited alternatively for each of four scanning sessions. Half of the participants started in room A and half started in room B. Each scanning session lasted ~11 min with a short break between the sessions, making a total functional scanning time of 50 min.

### 2.4 Behavioural analyses

For the pre-scan pointing task, we measured the mean 3D angular error for “within-room” trials and “across-room” trials. For the scanning direction judgment task, we first measured the overall accuracy (chance = 20%) to confirm whether participants knew their 3D direction in the virtual environment. We then tested whether participants were better at knowing their vertical or horizontal direction. In comparing vertical and horizontal performance, it was more informative to consider how much a participant’s response direction deviated from the true direction and not just whether they made a correct or wrong judgment. For example, when the true direction was 1 (“steep up”, Figure 2b), a participant could have selected either 2 (“shallow up”) or 4 (“shallow down”) and these errors were quantitatively different. To quantify the angular sensitivity, we defined the angular error of each trial by assigning 0° when participants chose the correct response; 30° when participants chose the adjacent direction such as 2 for 1, 60° when participants chose the direction 2 steps away from the correct direction such as 3 for 1, and so on. The mean angular error and response time (RT) was computed for vertical and horizontal questions respectively in each participant (excluding trials where participants did not respond within the time limit of 5 s, which occurred very rarely ‐ less than 1% of trials) and paired t-tests were used to compare the vertical and horizontal angular error and RT at the group level.

### 2.5 Scanning and pre-processing

T2*-weighted echo planar images (EPI) were acquired using a 3T Siemens Trio scanner (Siemens, Erlangen, Germany) with a 32-channel head coil. Scanning parameters optimised for reducing susceptibility-induced signal loss in areas near the orbitofrontal cortex and medial temporal lobe were used: 44 transverse slices angled at ‐30°, TR = 3.08 s, TE = 30 ms, resolution = 3 × 3 x 3 mm, matrix size = 64 x 74, z-shim gradient moment of ‐0.4mT/m ms (Weiskopf et al. 2006). Fieldmaps were acquired with a standard manufacturer’s double echo gradient echo field map sequence (short TE=10 ms, long TE=12.46 ms, 64 axial slices with 2 mm thickness and 1 mm gap yielding whole brain coverage; in-plane resolution 3 x 3 mm). After the functional scans, a 3D MDEFT structural scan was obtained with 1mm isotropic resolution.

Date were preprocessed using SPM12 (www.fil.ion.ucl.ac.uk/spm). The first five volumes from each functional session were discarded to allow for T1 equilibration effects. The remaining functional images were realigned to the first volume of each session and geometric distortion was corrected by the SPM unwarp function using the fieldmaps. Each participant’s anatomical image was then coregistered to the distortion corrected mean functional images. Functional images were normalised to MNI space.

### 2.6 fMRI analyses

#### 2.6.1 Delineating the anatomical regions of interest (ROIs)

We anatomically defined the ROIs – thalamus, EC, subiculum and RSC – that are known to contain HD cells. The thalamus ROI was extracted from the AAL atlas (Tzourio-Mazoyer et al., 2002). EC and subiculum ROIs were manually delineated on the group-averaged MRI scans from a previous independent study on 3D space representation (Kim et al., 2017) following the protocol in Pruessner et al., 2002. Although HD cells have been mainly found in presubiculum in animals, here we used a broader subiculum mask containing pre/parasubiculum because it was not feasible to distinguish these structures in our standard resolution fMRI images. The RSC ROI was also delineated on the group-averaged MRI scans. It contained Brodmann areas 29-30, located posterior to the splenium of corpus callosum (Vann et al., 2009). The number of functional voxels (3 x 3 x 3 mm) within each ROI (L = left, R = right) were as follows: thalamus_L, 302; thalamus_R, 286; RSC_L, 158; RSC_R, 135; EC_L, 47; EC_R, 49; subiculum_L, 34; subiculum_R, 34.

#### 2.6.2 Representational similarity analysis – ROIs

To examine whether each ROI contained vertical (pitch) or horizontal (azimuth) direction information or both, we used a multivoxel pattern analysis similar to that used in previous studies (e.g. Vass & Epstein, 2013; Carlin et al., 2011). This analysis compared the neural similarity measures to model similarity values predicted from multiple encoding hypotheses (which will be described in detail shortly). As a first step in the analysis, we estimated the neural responses to each 3D HD using a general linear model (GLM). The design matrix contained 25 main regressors which were boxcar functions that modelled the period when participants moved straight in one of 25 directions (5 levels for vertical pitch x 5 levels for horizontal azimuth), convolved with the SPM canonical hemodynamic response function. In addition, the occasional questions and blank screen periods (when participants came to the border of the spaceship) were separately modelled in the GLM as regressors of no interest. Six head realignment parameters were also included as nuisance regressors. The GLMs were applied for each scanning session in each participant.

We then computed the neural representational similarities between each direction using Pearson’s correlation using the multivoxel T-values within the ROIs that were estimated in the preceding GLM. We included all voxels within an ROI when calculating the multivoxel pattern similarities. Crucially, representational similarity was calculated between neural responses to the 3D directions when a participant was in different rooms of the virtual spaceship. This ensured that neural similarity was calculated between independent scanning sessions (because each room was alternatively visited in separate scanning sessions). More importantly, this across-room similarity analysis allowed us to detect relatively pure spatial direction information that was independent of view, which is naturally linked to HD. Figure 1b-e show example views when participants moved in two different directions in the two rooms. For instance, when we calculated the neural similarity between the “down-left” direction and “flat-right” direction, the correlation between “down-left” in room A (Figure 1b) and “flat-right” in room B (Figure 1e) and the correlation between “down-left” in room B (Figure 1c) and “flat-right” in room A (Figure 1d) were averaged. Therefore, the higher neural similarity between pairs of directions was not attributable to the higher visual similarity between the views associated with the directions within the same room. In summary, we calculated a symmetric 25 x 25 pairwise representational similarity matrix for each participant. We converted the similarity value (Pearson’s r) into a dissimilarity value by inverting it (1-r) for ease of later analysis.

Finally, these neural dissimilarity measures were compared to the vertical and horizontal directional encoding models using multiple regression. We used encoding models in which neural dissimilarity is linearly dependent on the difference in pitch or azimuth between two directions (Figure 3). For example, a vertical encoding model predicts that neural similarity between two directions that have the same pitch will be the highest, while neural similarity between two directions where pitch is ‐60° and 60° respectively will be the lowest, regardless of azimuth. We also included a visual texture similarity model to control for low-level visual similarity. Therefore, pitch distance, azimuth distance, visual similarity and a constant term were included in the multiple regression model. We computed visual texture similarity using the model of Renninger & Malik (2004). This visual control model was used in previous studies that investigated direction encoding (Vass & Epstein, 2013; Sulpizio et al., 2014; Kim et al., 2017).

**Figure 3.**
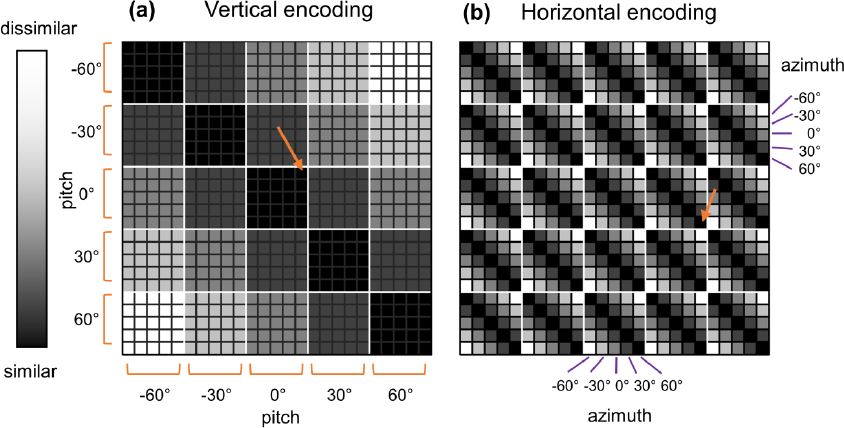
The model representational similarity matrix. The representational similarity matrix (25 x 25) contained pairwise similarity values between each of 25 unique 3D directions. (a) If the vertical direction was encoded, the neural similarity between the directions that share common vertical tilt, pitch would be high (dark colours), e.g. between (pitch, azimuth) = (0°,-60°) and (0°,60°) as indicated by the orange arrow. Similarity falls as the difference in pitch between two directions increases. (b) If the horizontal direction is encoded, the neural similarity between the directions that share a common horizontal angle, azimuth would be high (dark colours), e.g. between (pitch, azimuth) = (0°,60°) and (30°,60°), as indicated by the orange arrow. Similarity falls as the difference in azimuth between two directions increases.

Regression coefficients (beta) of each participant were fed into a group level analysis to test whether the neural response in the selected ROIs was explained by vertical or horizontal encoding models. We tested whether the regression coefficient was significantly greater than zero using a t-test. We also performed paired t-tests to compare the betas of the vertical and horizontal models to ascertain whether the neural response was more sensitive to one model or the other.

#### 2.6.3 Neural correlates of individual differences

We also tested whether there was a relationship between the direction information represented in the multivoxel pattern in our ROIs and behavioural performance during the scanning direction judgment task. For the behavioural performance measure, we used the mean angular error pooled across the vertical and horizontal direction questions given that the vertical and horizontal errors were highly correlated (Pearson’s r = 0.81, p < 0.001). We defined the direction information in individuals as the regression coefficient for the vertical and horizontal direction model in our ROIs. The Pearson correlation coefficient was used for the significance test.

#### 2.6.4 Representational similarity analysis – searchlight

While our main interest was in testing for the existence of vertical and horizontal direction information in our pre-specified ROIs, we also conducted a whole-brain searchlight analysis (Kriegeskorte et al., 2006) to test whether there was any other brain regions sensitive to vertical and horizontal direction. Moreover, the searchlight analysis complemented findings from the ROI analysis in the thalamus by providing additional anatomical localisation, given that the thalamus is a heterogeneous structure containing multiple functionally distinct nuclei. For localisation of thalamic structures, we relied on the WFUpickAtlas software (Lancaster et al. 1997; Lancaster et al. 2000; Maldjian et al., 2003) and a human thalamus atlas (Morel, 2007).

We performed the same representation similarity analysis using the multivoxel T-values within small spherical ROIs (radius 6mm) centred on each voxel across the whole brain. This generated regression coefficient maps for vertical and horizontal encoding models for each participant. These maps were fed into the group-level analysis (one-sample t-test) in SPM. We report voxel-wise p-values corrected for our anatomical ROIs. For the rest of the brain, we report voxels that survived whole-brain multiple comparison correction (family-wise error rate of 0.05). We used SPM voxel-level (peak-level) inference which computes corrected p-values using Random Field Theory.

## 3. Results

### 3.1 Behavioural results

The pre-scan pointing task involved participants wearing the VR head-mounted display and looking at the remembered position of balls while they were positioned at random locations. The group mean angular error was 21 ± 9° for within-room trials. Figure 1f shows an example view when a participant made a ~21° error, and we can see that the participant’s pointing direction (centre of the screen, a red crosshair) was reasonably close to the target ball. The error for across-room trials was slightly larger (28 ± 20°). This is unsurprising, because participants had to orient themselves to the target ball behind the wall. Given this overall good level of performance, we are confident that participants went into the subsequent scanning experiment with a reasonable sense of orientation in the 3D virtual environment.

During scanning, participants were moved in a preprogrammed 3D trajectory and were occasionally asked about their movement direction, either vertically or horizontally. The mean accuracy (74 ± 16%) was well above chance level (20%), suggesting that participants were able to keep track of their movement direction. We found that participants made significantly smaller errors for the vertical questions compared to the horizontal questions (t(29) = ‐2.43, p = 0.021, Figure 2c). We also observed a small, but significant, difference in RT in favour of the horizontal questions (vertical = 1.79 ± 0.36 s, horizontal = 1.67 ± 0.38 s, t(29) = 2.57, p = 0.015).

### 3.2 fMRI results – ROIs

We investigated whether multivoxel patterns in our ROIs contained vertical and/or horizontal direction information. The right RSC showed both vertical and horizontal direction information (vertical, t(29) = 3.69, p = 0.001; horizontal, t(29) = 2.05, p = 0.050, Figure 4a), but this region was significantly more sensitive to vertical direction (paired t-test, t(29) = 2.61, p = 0.014). In contrast, the left thalamus showed only horizontal direction encoding (t(29) = 2.81, p = 0.009, Figure 4b), and horizontal encoding was significantly stronger than vertical encoding (paired t-test, t(29) = ‐2.36, p = 0.025). The right thalamus and left subiculum also showed horizontal direction information (thalamus, t(29) = 2.27, p = 0.031; subiculum, t(29) = 2.63, p = 0.013, Figure 4c,d), but direct comparison between vertical and horizontal sensitivity was not significant. Bilateral EC, right subiculum, and left RSC did not show any significant evidence of vertical or horizontal direction encoding.

**Figure 4.**
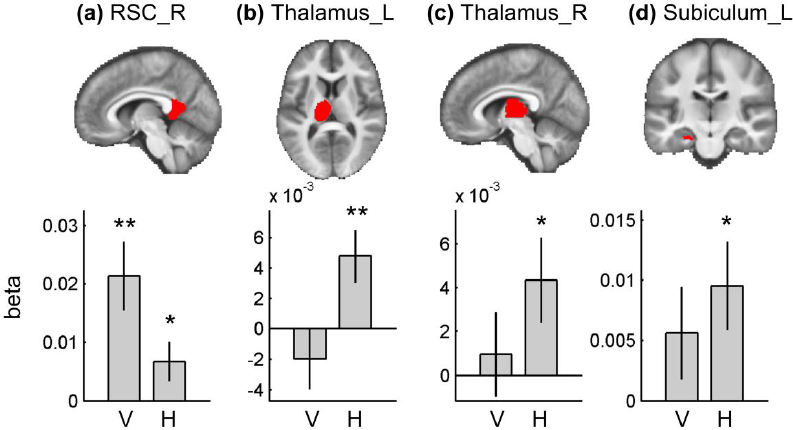
Multivoxel pattern analysis in the ROIs. Each ROI is overlaid on the group averaged structural MR image on the top row. (a) Right RSC showed both vertical and horizontal direction encoding, but it was more sensitive to vertical direction. Bilateral thalamus (b, c) and left subiculum (d) showed only horizontal direction encoding. V, vertical; H, horizontal; R, right; L, left. Error bars are SEM. ** p<0.01, * p<0.05.

### 3.3 Individual differences

The above analysis revealed evidence of vertical and horizontal direction information in RSC, thalamus and subiculum at the group level. We then tested whether direction information in these regions could explain the individual differences in behavioural performance during our direction judgment test. We found that vertical direction information in the right RSC was significantly correlated with angular error (r = −0.45, n = 30, p = 0.009, Figure 5). This means that participants whose right RSC showed more vertical direction information were more accurate at making direction judgments. Horizontal direction information in the right RSC, bilateral thalamus, and left subiculum was not correlated with behaviour (p>0.05).

**Figure 5.**
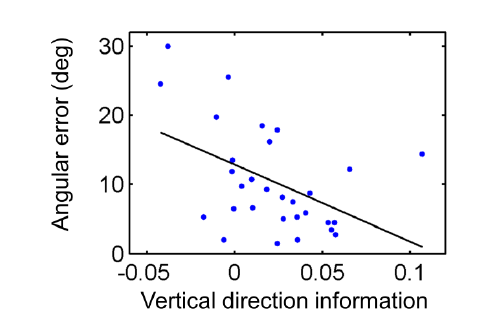
Participants whose right RSC exhibited more vertical direction information were better at the direction judgment task (i.e. had a smaller angular error); r = ‐0.45, n = 30, p = 0.009.

### 3.4 fMRI results – searchlight

A whole-brain searchlight analysis for vertical direction encoding identified bilateral RSC (right, peak at [9, −58, 8], t(29) = 5.62, p = 0.001, 16 voxels with T>4.058 within RSC; left, [−9, −46, 2], t(29) = 5.04, p = 0.003, 6 voxels with T>4.058 within RSC; small volume corrected for bilateral RSC masks, Figure 6a), similar to the finding from the ROI analysis. Clusters in lingual gyrus (peak, [−12, −61, 2], t(29) = 7.29, p = 0.002, 10 voxels with T>6.02; [3, −61, 8], t(29) = 6.87, p = 0.005, 5 voxels with T>6.02) and cuneus (peak, [6,−82,17], t(29) = 7.24, p = 0.002, 13 voxels with T>6.02) also showed vertical direction information.

**Figure 6.**
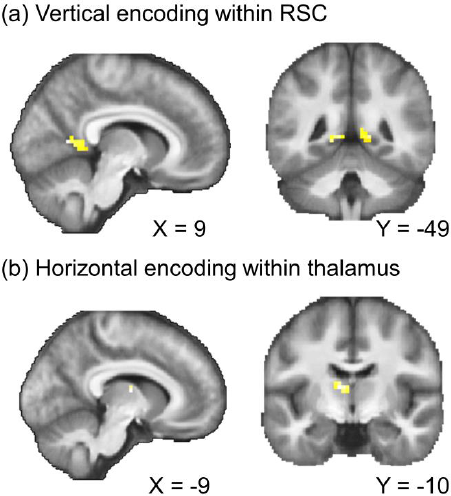
Searchlight results. (a) Vertical direction information within the bilateral RSC mask. (b) Horizontal direction information within the bilateral thalamus mask; p<0.001 uncorrected for display purposes. See the main text for the other regions that survived whole-brain multiple comparison correction.

Horizontal direction information was observed in the anterior part of the left thalamus (peak at [−9, −10, 11], t(29) = 4.73, p = 0.016, 3 voxels with T>4.313; small volume corrected for bilateral thalamus masks, Figure 6b). The peak coordinate is most likely located in the ventral anterior nucleus, but we caveat this localisation by noting that the spatial resolution of our fMRI scans (3 mm) was not fine enough to identify small thalamic nuclei with confidence. Furthermore, neural responses in the neighbouring thalamic nuclei could have contributed to this finding due to the nature of the multivoxel pattern analysis (6 mm radius). We also observed a voxel in the left subiculum which showed horizontal direction information, as in the earlier ROI analysis ([−27, −25, −16], t(29) = 3.58, p = 0.039, 1 voxel with T >3.483; small volume corrected for the bilateral subiculum mask). At the whole-brain corrected level, horizontal direction information was also observed in the central sulcus ([−33, −22, 50], t(29) = 8.63, p<0.001, 6 voxels with T>6.02), supplementary motor cortex ([-6, 5, 53], t(29) = 6.10, p = 0.041, 1 voxel with T>6.02), and visual cortex ([−9, −82, −10], t(29) = 6.22, p = 0.029, 1 voxel with T>6.02; [−9, −79, 5], t(29) = 6.04, p = 0.047, 1 voxel with T>6.02; [-6,-73,-7], t(29) = 6.24, p = 0.027, 1 voxel with T>6.02). Of note, the number of voxels in a searchlight result should be interpreted with caution. Even when a single voxel is reported as significant, this result is driven by multiple neighbouring voxels (6 mm radius) due to the nature of searchlight analyses. Furthermore, our data had a relatively small amount of smoothness compared to typical fMRI studies because we did not apply spatial smoothing during preprocessing or the group-level analysis in order to retain the best spatial specificity.

## 4. Discussion

In this study we investigated how 3D HD was encoded in the human brain when participants moved in a volumetric space. Using a virtual reality environment and fMRI multivoxel pattern similarity analysis, we found that the thalamus and subiculum were sensitive to the horizontal component of 3D HD. By contrast, vertical heading information was dominant in RSC, and vertical direction information in RSC was significantly correlated with behavioural performance during a direction judgment task.

The anterior thalamic nuclei (ATN) are important subcortical structures for spatial navigation and memory (Jankowski et al., 2013). Within the hierarchy of the HD cell network, the ATN receive vestibular inputs via the lateral mammillary nuclei and project to higher cortical areas including RSC and dorsal presubiculum (Taube, 2007). Most HD cells in the ATN have been recorded when rodents move on a 2D plane. A previous human fMRI study also found 2D direction information in the thalamus (Shine et al., 2016). The current study, therefore, extends our understanding of the HD system by providing the first evidence that the thalamus (especially the anterior portion) encodes horizontal heading even when participants move in a volumetric 3D space. The lack of vertical direction information in the thalamus resembles the early finding of HD cells in the lateral mammillary nuclei, which were insensitive to the vertical head tilt of rats (Stackman & Taube 1998), although we should be mindful of the difference in structures (thalamus versus mammillary nuclei) and environments (3D spaceship versus 2D plane), and the limitations of the recording apparatus used in this early rat study. The vertical insensitivity of the thalamus might also be related to previous findings that showed HD cells in the rat ATN maintained the preferred direction on the vertical wall as if the wall was an extension of the floor, and the HD cells only cared about the rotation along the body axis, not the rotation of the body axis relative to the vertical gravity axis (Calton & Taube, 2005; Taube et al., 2013).

Why the thalamus was not sensitive to vertical pitch is an interesting question that requires further investigation. One possible explanation is that the vestibular system, which is responsible for angular integration and updating of the responses of HD cells in the thalamus, might be less sensitive to vertical rotation because humans are surface-based animals and we infrequently rotate vertically. Although our participants’ heads were immobilised during scanning, vestibular inputs they experienced during the pre-scan task with the VR head-mounted display might have been reinstated by visual cues during scanning and contributed to HD encoding, as suggested by a previous study (Shine et al. 2016). Furthermore, optic flow during scanning could have stimulated the vestibular nuclei (Glasauer, 2005), and indeed HD cells in the thalamus of rats have been found to be modulated by pure optic flow without visual landmarks (Arleo et al., 2013). Vertical and horizontal optokinetic responses are known to activate both common and unique vestibular nuclei (Bense et al., 2006).

It is also possible that vertical information might be more evident in the thalamus if we studied navigation in a real environment instead of a virtual environment. Recently, Laurens et al. (2016) found cells tuned to gravity (vertical tilt) in the macaque anterior thalamus using a rotatable apparatus (Laurens et al., 2016). Even though the pre-scan immersive training and the optic flow during our scanning experiment could have enhanced the HD signal, physical head tilt and acceleration was missing in our fMRI study. Given the importance of vestibular inputs in generating and maintaining stable HD signals, as shown by lesion studies in animals (e.g. Muir et al., 2009; Yoder & Taube, 2009), 3D HD encoding should be studied in freely moving participants in the future.

Our next finding concerns the subiculum. The presubiculum is reciprocally connected to the anterior thalamus, and a lesion in the thalamus disrupts HD cells in the presubiculum (Goodridge & Taube, 1997). To date, presubiculum is the only brain structure where HD cells have been recorded in animals exploring a volumetric space (Finkelstein et al., 2015). In this bat study, cells that were sensitive to either horizontal only or vertical only heading as well as conjunctive cells were found in presubiculum. In the present study, we found only horizontal direction information in the human subiculum. This might be attributable to a difference in species (bat, a flying animal, versus human surface-dwellers) or to methodological differences. Unlike invasive recordings, fMRI measures aggregate neural responses. Therefore, if the human subiculum contains more azimuth-tuned cells than pitch-tuned cells, similar to bats (Finkelstein et al., 2015), azimuth information might be more easily detected by fMRI. The existence of azimuth and pitch encoding in the subiculum would be better addressed in a future fMRI study with higher spatial resolution, if indeed a similar anatomical gradient of azimuth, pitch and conjunctive cells also exists in the human brain (Finkelstein et al., 2015).

Unlike the thalamus or subiculum, the right RSC showed vertical direction information, although horizontal information was also present in this region. Therefore, in principle it seems that RSC could serve as a 3D compass on its own. Our finding of a significant correlation between vertical direction information in the RSC and behavioural accuracy might reflect the functional relevance of RSC for processing 3D direction information (although it is unclear why only vertical direction information and not horizontal direction information in this region correlated with individual differences). The dominance of vertical information in the RSC was concordant with our previous finding of vertical direction encoding when participants moved on a 3D rollercoaster (Kim et al., 2017). One explanation could be that visual cues might be more salient for the vertical axis compared to the horizontal axis. Within the HD system, RSC is directly connected to early visual cortex (Kobayashi & Amaral, 2003) and HD cells in RSC are dominated by local visual landmarks (Jacob et al., 2017). Of note, presubiculum is also known to have direct connections with V2 in rodents (Vogt & Miller, 1983), but we are not aware of direct connections between the presubiculum and early visual cortex in primates.

Behaviourally, participants were more accurate at judging vertical direction, and some participants anecdotally reported that they felt the vertical direction judgment was easier (note, however, that the RT was longer) because of the views of the ceiling and floor, even though we also designed the side walls to provide clear polarisation cues for the horizontal direction. Views are naturally dependent on HD, and the horizontal component of HD has less influence on views as the vertical tilt increases in 3D space. For example, if a participant looks straight towards East or West (zero vertical tilt), the views can be very different due to distinct landmarks. In contrast, when the vertical tilt is 90°, the participant looks straight up in the sky and the views will be similar regardless of whether they face East or West. Although we tried to orthogonalise the view and HD by measuring the neural similarity between pairs of directional responses across different rooms in our virtual environment (as we explained in the Materials and Methods), and we also added the low-level visual texture similarity regressor for extra control, there still remains a possibility that the views were more similar when the vertical tilts were similar compared to when the horizontal direction was similar. This could reflect the nature of the relationship between HD and view in 3D space, rather than being a particular feature of our virtual environment.

Related to the vertical-horizontal asymmetry, one interesting question is the potential influence of an explicit cognitive task on the neural representation of HD. In the current experiment, we occasionally asked participants to indicate their vertical or horizontal direction between movements. This task could be answered rapidly and easily, thus minimising interruption to movement and eschewing the need for additional scanning time, while ensuring that participants paid attention to their 3D movement direction. However, the explicit and separate questions for vertical and horizontal directions might have contributed to the encoding of vertical and horizontal information in different brain regions. Vertical and horizontal information might be more homogenously represented in these brain regions if participants move freely in 3D space without explicitly paying attention to the vertical and horizontal components of direction. Experimenters could then avoid using the terms “vertical” and “horizontal” during the experiment, and participants could be asked to directly indicate their 3D direction (although we note that it is almost impossible to indicate precisely and rapidly one’s 3D direction without dividing it to vertical and horizontal components). Alternatively, cognitive tasks that test an explicit awareness of movement direction could be removed, given that HD cells are often recorded in rodents when animals forage in an environment without active navigation or a spatial memory test.

In contrast, more spatially demanding tasks, such as 3D path integration with multiple pitch, roll and yaw rotations (Vidal et al., 2004), might result in stronger HD signals both vertically and horizontally. Different behavioural paradigms, where some are more explicit than others, should be utilised to study 3D HD encoding in the future. Nevertheless, we believe that studying vertical and horizontal components will remain pertinent to the research field of 3D spatial encoding regardless of behavioural paradigms, because all species on earth are under the influence of gravity which distinguishes the vertical from the horizontal axis. Even astronauts in microgravity have reported that they tend to orient themselves to local surfaces and use the words “up” and “down” (Oman, 2007).

In summary, the current study presented the first evidence showing that thalamus, subiculum and RSC – the ‘classic’ HD system that has been identified when tested on a horizontal 2D plane – also encodes vertical and horizontal heading in 3D space. We suggest that these brain structures play complementary roles in processing 3D direction information regarding angular integration and visual cues. Future studies of the HD system in real volumetric space should elucidate specifically how each sensory modality (visual, vestibular, proprioceptive) and physical gravity contributes to HD encoding in these brain structures. This could, perhaps, be facilitated by using the recently-developed ‘mobile’ magnetoencephalography brain scanner which allows head movements while measuring neural activity in humans, including from deep brain structures such as those implicated in the head direction system (Boto et al., 2018).

## ACKNOWLEDGEMENTS

The authors thank Christian Lambert and Marshall Dalton for their advice on thalamic and hippocampal anatomy.

The authors declare no competing financial interests.

## SUPPORTING INFORMATION

**SUPPORING FIGURE S1.**
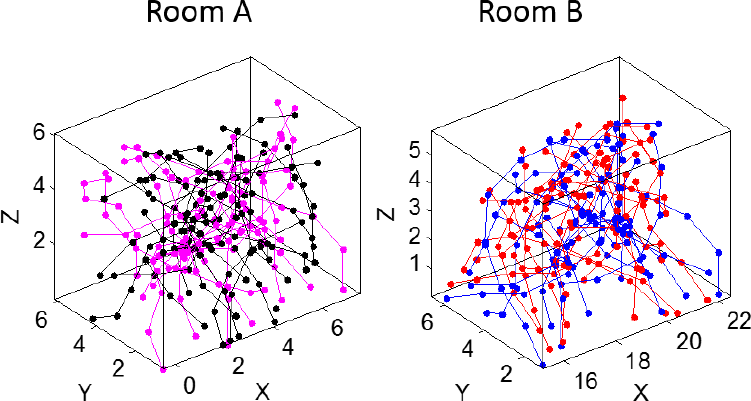
3D movement trajectories during scanning. Different colors represent different scanning sessions.

## REFERENCES

Arleo, A., Dejean, C., Allegraud, P., Khamassi, M., Zugaro, M. B., & Wiener, S. I. (2013). Optic flow stimuli update anterodorsal thalamus head direction neuronal activity in rats. Journal of Neuroscience, 33(42), 16790–16795.

Baumann, O., & Mattingley, J. B. (2010). Medial parietal cortex encodes perceived heading direction in humans. Journal of Neuroscience, 30(39), 12897–901.

Bense, S., Janusch, B., Vucurevic, G., Bauermann, T., Schlindwein, P., Brandt, T., … Dieterich, M. (2006). Brainstem and cerebellar fMRI-activation during horizontal and vertical optokinetic stimulation. Experimental Brain Research, 174(2), 312–323.

Boto, E., Holmes, N., Leggett, J., Roberts, G., Shah, V., Meyer, S. S., … Brookes, M. J. (2018). Moving magnetoencephalography towards real-world applications with a wearable system. Nature, 555(7698), 657–661.

Burak, Y., & Fiete, I. R. (2009). Accurate path integration in continuous attractor network models of grid cells. PLoS Computational Biology, 5(2). e100291.

Calton, J. L., Stackman, R. W., Goodridge, J. P., Archey, W. B., Dudchenko, P. A., & Taube, J. S. (2003). Hippocampal place cell instability after lesions of the head direction cell network. Journal of Neuroscience, 23(30), 9719–9731.

Calton, J. L., & Taube, J. S. (2005). Degradation of head direction cell activity during inverted locomotion. Journal of Neuroscience, 25(9), 2420–2428.

Carlin, J. D., Calder, A. J., Kriegeskorte, N., Nili, H., & Rowe, J. B. (2011). A head view-invariant representation of gaze direction in anterior superior temporal sulcus. Current Biology, 21(21), 1817–1821.

Chadwick, M. J., Jolly, A. E. J., Amos, D. P., Hassabis, D., & Spiers, H. J. (2015). A goal direction signal in the human entorhinal/subicular region. Current Biology, 25(1), 87–92.

Cullen, K. E., & Taube, J. S. (2017). Our sense of direction: Progress, controversies and challenges. Nature Neuroscience, 20(11), 1465–1473.

Finkelstein, A., Derdikman, D., Rubin, A., Foerster, J. N., Las, L., & Ulanovsky, N. (2015). Three-dimensional head-direction coding in the bat brain. Nature, 517(7533), 159–164.

Glasauer, S. (2005). Vestibular and motor processing for head direction signals. In S. I. Wiener & J. S. Taube (Eds.), Head Direction Cells and the Neural Mechanisms of Spatial Orientation (pp. 113–135). Cambridge, MA: The MIT press.

Goodridge, J. P., & Taube, J. S. (1997). Interaction between the postsubiculum and anterior thalamus in the generation of head direction cell activity. Journal of Neuroscience, 17(23), 9315–9330.

Indovina, I., Maffei, V., Pauwels, K., Macaluso, E., Orban, G. A., & Lacquaniti, F. (2013). Simulated self-motion in a visual gravity field: Sensitivity to vertical and horizontal heading in the human brain. NeuroImage, 71, 114–124.

Indovina, I., Maffei, V., Mazzarella, E., Sulpizio, V., Galati, G., & Lacquaniti, F. (2016). Path integration in 3D from visual motion cues: A human fMRI study. NeuroImage, 142, 512–521.

Jankowski, M. M., Ronnqvist, K. C., Tsanov, M., Vann, S. D., Wright, N. F., Erichsen, J. T., & O’Mara, S. M. (2013). The anterior thalamus provides a subcortical circuit supporting memory and spatial navigation. Frontiers in Systems Neuroscience, 7, 45.

Kim, M., Jeffery, K. J., & Maguire, E. A. (2017). Multivoxel pattern analysis reveals 3D place information in the human hippocampus. The Journal of Neuroscience, 37(16), 2703–16.

Kobayashi, Y., & Amaral, D. G. (2003). Macaque monkey retrosplenial cortex: II. Cortical afferents. Journal of Comparative Neurology, 466(1), 48–79.

Kriegeskorte, N., Goebel, R., & Bandettini, P. (2006). Information-based functional brain mapping. Proceedings of the National Academy of Sciences, 103(10), 3863–3868

Lancaster, J. L., Rainey, L. H., Summerlin, J. L., Freitas, C. S., Fox, P. T., Evans, A. C., … Mazziotta, J. C. (1997). Automated labeling of the human brain: A preliminary report on the development and evaluation of a forward-transform method. Human Brain Mapping, 5(4), 238–242.

Lancaster, J. L., Woldorff, M. G., Parsons, L. M., Liotti, M., Freitas, C. S., Rainey, L., … Fox, P. T. (2000). Automated Tailairach atlas labels for functional brain mapping. Human Brain Mapping, 10, 120–131.

Laurens, J., Kim, B., Dickman, J. D., & Angelaki, D. E. (2016). Gravity orientation tuning in macaque anterior thalamus. Nature Neuroscience, 19(12), 1566–1568.

Maldjian, J. A., Laurienti, P. J., Kraft, R. A., & Burdette, J. H. (2003). An automated method for neuroanatomic and cytoarchitectonic atlas-based interrogation of fMRI data sets. NeuroImage, 19(3), 1233–1239.

Marchette, S., Vass, L., Ryan, J., & Epstein, R. (2014). Anchoring the neural compass: coding of local spatial reference frames in human medial parietal lobe. Nature Neuroscience, 17(11), 1598–1606.

Morel, A. (2007). Stereotactic atlas of the human thalamus and basal ganglia. New York: Informa Healthcare USA.

Morey, R. D. (2008). Confidence intervals from normalized data: A correction to Cousineau (2005). Tutorials in Quantitative Methods for Psychology, 4(2), 61–64.

Muir, G.M., Brown, J.E., Carey, J.P., Hirvonen, T.P., Della Santina, C.C., Minor, L.B., Taube, J.S. (2009). Disruption of the head direction cell signal after occlusion of the semicircular canals in the freely moving Chinchilla. Journal of Neuroscience, 29, 14521–14533.

Oman C. (2007) Spatial orientation and navigation in microgravity. In: Mast F., Jäncke L. (Eds) Spatial Processing in Navigation, Imagery and Perception. Springer, Boston, MA

Page, H. J. I., Wilson, J. J., & Jeffery, K. J. (2018). A dual-axis rotation rule for updating the head direction cell reference frame during movement in three dimensions. Journal of Neurophysiology, 119, 192–208.

Pruessner, J. C., Köhler, S., Crane, J., Pruessner, M., Lord, C., Byrne, A., … Evans, A. C. (2002). Volumetry of temporopolar, perirhinal, entorhinal and parahippocampal cortex from high-resolution MR images: considering the variability of the collateral sulcus. Cerebral Cortex, 12(12), 1342–1353.

Raudies, F., Brandon, M.P., Chapman, G.W., Hasselmo, M.E. (2015). Head direction is coded more strongly than movement direction in a population of entorhinal neurons. Brain Research, 1621, 355–367.

Renninger, L. W., & Malik, J. (2004). When is scene identification just texture recognition? Vision Research, 44(19), 2301–11.

Robertson, R. G., Rolls, E. T., Georges-François, P., & Panzeri, S. (1998). Head direction cells in the primate hippocampal formation. Hippocampus, 9, 206–219.

Shine, J. P., Valdes-Herrera, J. P., Hegarty, M., & Wolbers, T. (2016). The human retrosplenial cortex and thalamus code head direction in a global reference frame. Journal of Neuroscience, 36(24), 6371–6381.

Stackman, R., & Taube, J. (1998). Firing properties of rat lateral mammillary single units: head direction, head pitch, and angular head velocity. The Journal of Neuroscience, 18(21), 9020–9037.

Sulpizio, V., Committeri, G., & Galati, G. (2014). Distributed cognitive maps reflecting real distances between places and views in the human brain. Frontiers in Human Neuroscience, 8, 716.

Taube, J. S., Stackman, R. W., Calton, J. L., & Oman, C. M. (2004). Rat head direction cell responses in zero-gravity parabolic flight. Journal of Neurophysiology, 92(5), 2887–2997.

Taube, J. S. (2007). The head direction signal: origins and sensory-motor integration. Annual Review of Neuroscience, 30(1), 181–207.

Taube, J. S., Wang, S. S., Kim, S. Y., & Frohardt, R. J. (2013). Updating of the spatial reference frame of head direction cells in response to locomotion in the vertical plane. Journal of Neurophysiology, 109(3), 873–888.

Tzourio-Mazoyer, N., Landeau, B., Papathanassiou, D., Crivello, F., Etard, O., Delcroix, N., & Joliot, M. (2002). Automated anatomical labeling of activations in SPM using a macroscopic anatomical parcellation of the MNI MRI single-subject brain. NeuroImage, 15(1), 273–289.

Vann S.D., Aggleton J.P., Maguire EA. (2009). What does the retrosplenial cortex do? Nature Reviews Neuroscience, 10, 792–802.

Vass, L. K., & Epstein, R. A. (2013). Abstract representations of location and facing direction in the human brain. Journal of Neuroscience, 33(14), 6133–6142.

Vidal, M., Amorim, M.-A. A., & Berthoz, A. (2004). Navigating in a virtual three-dimensional maze: how do egocentric and allocentric reference frames interact? Cognitive Brain Research, 19(3), 244–258.

Vogt, B. A., & Miller, M. W. (1983). Cortical connections between rat cingulate cortex and visual, motor, and postsubicular cortices. Journal of Comparative Neurology, 216(2), 192–210.

Weiskopf, N., Hutton, C., Josephs, O., & Deichmann, R. (2006). Optimal EPI parameters for reduction of susceptibility-induced BOLD sensitivity losses: a whole-brain analysis at 3 T and 1.5 T. NeuroImage, 33(2), 493–504.

Yoder, R.M., Taube, J.S. (2009). Head direction cell activity in mice: robust directional signal depends on intact otolith organs. Journal of Neuroscience, 29, 1061–1076.

